# Exo70 promotes herpesvirus secretory vesicle tethering and virion release in an exocyst complex-dependent manner

**DOI:** 10.1101/2024.03.08.584044

**Authors:** Wenqing Ma, Yanan Xu, Jialin Xie, Luteng He, Lin Wang, Jie Yu, Fachao Sun, Daniel Chang He, Hongmei Wang, Hongbin He

## Abstract

The exocyst complex, a multisubunit protein complex, mediates secretory vesicles fusion with the plasma membrane (PM) to deliver materials to the cell surface or to release cargoes to the extracellular space, but whether it is related to viral secretory vesicles tethering and virion release is unknown. Here, we identified that the exocyst complex subunit Exo70 promotes the trafficking of herpesvirus secretory vesicles and the release of mature virions in an exocyst complex-dependent manner. Mechanistically, mutation of Exo70 Lys632 and Lys635 diminishes viral secretory vesicles anchoring to PM. Moreover, the small GTPase Rab11a is necessary for the transport of viral secretory vesicles, and the Snapin-Exo70-SNAP23 axis is involved in fusion of viral secretory vesicles to the PM and release of virions. Most significantly, we discovered that Endosidin2 (ES2), an inhibitor of exocytosis via the exocyst complex, provides protection against herpesvirus infection in cells and mice. Overall, these findings unveil a previously uncharacterized role and mechanism of the exocyst complex in viral replication, highlighting its potential as an effective strategy against herpesvirus infection.

**Author summary:** Herpesviruses, such as herpes simplex virus 1 (HSV-1) and bovine herpes virus type 1 (BoHV-1), are highly prevalent pathogens that establish lifelong infections and cause various diseases in humans and animals. While the mechanisms underlying the transport of herpesvirus secretory vesicles from the Golgi to the PM and the subsequent viral release are still poorly understood. In our study, we have discovered that Exo70 promotes the trafficking of HSV-1 and BoHV-1 secretory vesicles and facilitates the release of mature virions in an exocyst complex-dependent manner. Mechanistically, both knockdown of Exo70 and a phosphatidylinositol 4,5-bisphosphate (PI(4,5)P2)-binding-deficient mutant of Exo70 caused virions to be trapped in the cytoplasm and unable to tether to PM, indicating that the binding of Exo70 to PI(4,5)P2 is critical for the docking virus secretory vesicles with PM. Additionally, the Snapin-Exo70-SNAP23 axis plays a pivotal role in the fusion between viral secretory vesicles and the PM, ultimately facilitating the release of virions. We found that Rab11a is necessary for viral secretory vesicle trafficking mediated by Exo70. Most importantly, we discovered that ES2, a transport inhibitor that combines with Exo70 to terminate the final stage of exocytosis via the exocyst complex, can protect cells and mice from α-herpesvirus infection.

## Introduction

Herpesviruses are common pathogens capable of establishing lifelong infections and causing a variety of diseases [1]. It consists of four morphologically distinct components: the core, capsid, tegument, and envelope [2–5]. The lytic replication cycle of the herpesvirus includes entry into host cells, viral genome uncoating, gene replication, capsid assembly and egress from the nucleus, maturation and envelopment of viral particles in the cytoplasm and exocytosis of mature virions [6, 7]. Several studies have presented evidence supporting the trans-Golgi network (TGN) as the site of secondary envelopment for herpesviruses [8–11]. However, our understanding of how mature herpesviruses are transported and released from cells following secondary envelopment is still limited [12].

The exocyst is an octameric protein complex that is involved in maintaining cell morphology, building cell connections and promoting cell migration [13]. The exocyst complex consists of the subunits Sec3, Sec5, Sec6, Sec8, Sec10, Sec15, Exo70 and Exo84 [14]. Among these eight subunits, Sec3, Sec5, Sec6, and Sec8 assemble into a tight quaternary subcomplex, whereas Sec10, Sec15, Exo70, and Exo84 form a separate subcomplex [15, 16]. Exo70 protein, a fundamental component of the evolutionarily conserved octameric exocyst complex, is responsible for tethering post-Golgi vesicles to the plasma membrane (PM) in preparation for SNARE-mediated membrane fusion [17]. Following vesicle tethering, the physical association of Exo70 with phosphatidylinositol 4,5-bisphosphate (PI(4,5)P2) may act as an ’anchor’ that attaches the exocyst complex and vesicles to the PM [18]. While the exocyst and Exo70 subunit facilitate the tethering of post-Golgi secretory vesicles to the PM and assist in cargo egress, their role in regulating virus release remains unknown.

The fusion of vesicles with the PM is a crucial step in the cargo egress. Soluble N-ethylmaleimide-sensitive factor attachment protein receptor (SNARE) proteins mediate membrane fusion events in eukaryotic cells [19]. SNAREs are members of a family of proteins distributed into two groups: SNAREs in one set are attached to the vesicle (v-SNAREs), and those in a complementary set are attached to the vesicle’s target membrane (t-SNAREs) [20, 21]. The fusion reaction is initiated when the v-SNAREs with the t-SNAREs to form a trans-SNARE complex [22]. The trans-SNARE complex is essential for vesicle fusion because SNARE proteins provide the energy for membrane fusion to deliver cargos [23]. Exo70 may participate in the assembly of trans-SNARE complex. Synaptosome-associated protein 23 kDa (SNAP23) is a t-SNARE that is widely expressed in the cell membrane. Exo70 indirectly interacts with SNAP23 by sequentially binding Snapin, mediating the fusion of Glut4-secreting vesicles and PM [24]. Nevertheless, it remains unclear whether the interaction between Exo70, Snapin, and SNAP23 plays a role in the release of herpesvirus particles.

The exocyst complex cannot directly bind to secretory vesicles, and therefore, the assembly process requires the assistance of GTPase [25]. GTPase Rab11 acts in conjunction with exocyst complex to mediate the steps in cargo transport [26]. During E. coli clearance, the exocyst complex subunit Sec6 recruits and activates Rab11a, facilitating bacterial transport out of the cell with the assistance of dynein [27]. Nonetheless, the potential cooperation between Rab11a and Exo70 in the transportation of virions has not been documented.

Our study revealed that Exo70, in a manner dependent on the exocyst complex, facilitates the transport of herpesvirus secretory vesicles via Rab11a from the TGN, anchoring them to the PM by binding to PI(4,5)P2 and enabling the release of virions through the Snapin-Exo70-SNAP23 axis. Importantly, our findings revealed that ES2, an inhibitor of exocytosis mediated by the exocyst complex, has the ability to effectively suppress viral replication.

## Results

### Herpesvirus secretory vesicles are transported to the PM via the Golgi-PM transport pathway

In the final stages of herpesvirus replication, a viral nucleocapsid buds into a vesicle of TGN/endosome origin, acquiring an envelope and an outer vesicular membrane. The virus-containing secretory vesicle then traffics to the PM, to which it fuses, exposing the mature virion [28]. In this study, we utilized transmission electron microscopy to observe HSV-1 and BoHV-1 in infected cells. Circular viral particles with a diameter of approximately 150 nm were exclusively found on the Golgi body, not in uninfected cells. And the virions on the Golgi apparatus were coated with two layers of envelope (Fig 1A and S1A). This indicates that the secondary envelope of herpesvirus can occur on the Golgi apparatus. After blocking the Golgi-PM transport pathway with monensin treatment, the colocalization of the BoHV-1 gB protein and GM130 (a Golgi marker) increased significantly, and gB was mainly distributed in the Golgi body with less localization on PM, while in the monensin-untreated control group, the colocalization of the gB protein and GM130 was reduced, and the gB protein was distributed in the cytoplasm and PM (Fig 1B, 1C, and 1D). Comparable outcomes were observed in HSV-1-infected cells (Fig S1B, S1C, and S1D). These results indicated that monensin treatment blocked the transport of herpesvirus to the PM, which was also confirmed by transmission electron microscopy. The number of virions adjacent to the Golgi apparatus in BoHV-1-infected MDBK cells treated with monensin was significantly higher than that in the control group (indicated by the black arrow) (Fig 1E), while the number of virions released into the extracellular space was significantly lower than that in the control group (Fig 1F and 1G). Similarly, in Vero cells infected with HSV-1, the number of virions near the Golgi bodies following monensin treatment was greater than that in the DMSO treatment group (Fig S1E), while the number of virions released into the extracellular space was lower (Fig S1F and S1G). These compelling results emphasize the significant role of the Golgi-PM transport pathway in herpesvirus transport to the PM.

**Fig 1.**
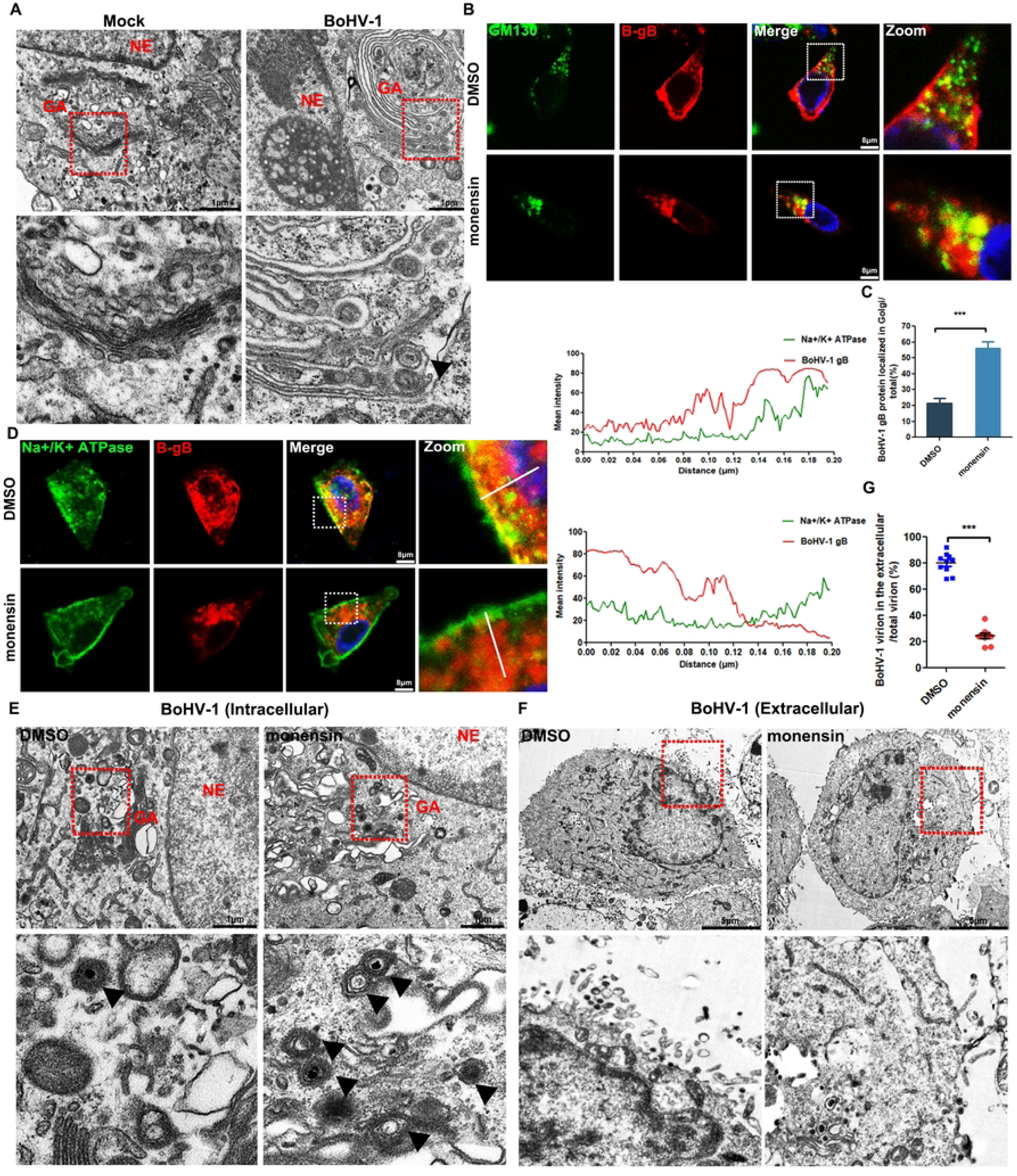
Virions were trapped on the Golgi apparatus by blocking the Golgi-PM transport pathway. (A) Transmission electron microscopy showing the secondary envelope of the virions in Vero cells, which infected with BoHV-1 (MOI=0.1) for 24 h and analyzed by transmission electron microscopy. NE, nucleus; GA, Golgi apparatus. Scale bar, 1 μm. (B) Confocal microscopy images of MDBK cells treated with DMSO or monensin for 4 h and subsequently infected with BoHV-1 (MOI=1) for 12 h. The cells were fixed and stained for GM130 (green), gB (red), and chromatin (Hoechst, blue). Scale bar, 8 μm. (C) The percentages of gB protein localized in the Golgi were quantified. (D) Immunofluorescence of Vero cells treated with DMSO or monensin, infected with BoHV-1, fixed and stained for Na^+^/K^+^ ATPase (green) and gB (red). The fluorescence intensity profiles of Na^+^/K^+^ ATPase (green) and gB (red) were measured along the line drawn by ImageJ. (E) Transmission electron microscopy showing the virions adjacent to the GA. Arrowheads indicate virions. (F) Transmission electron microscopy showing the extracellular virions. Scale bar, 5 μm. (G) The percentages of extracellular virions were quantified. Data are presented as the means ± SEMs of three independent experiments.

### Silencing subunits of the exocyst complex inhibited herpesvirus replication

The exocyst complex consists of eight highly conserved subunits. Every four subunits form a subcomplex: Sec3, Sec5, Sec6, and Sec8 form subcomplex I, and Sec10, Sec15, Exo70, and Exo84 form subcomplex II (Fig 2A). The assembly of the two subcomplexes at the appropriate time and location forms the complete octameric complex and ensures the spatiotemporal specificity of a vesicle-tethering event [29, 30]. We selected and separately silenced two subunits from each subcomplex to investigate the functions of exocyst complex subunits in herpesvirus replication. Sec3, Sec6, Exo70, and Exo84 were silenced in Vero cells, which were then infected with HSV-1. 24 h later, the supernatant was collected to detect viral titers. The results showed that the silencing of four subunits resulted in the decrease of viral titers in the supernatant to varying degrees, with silencing of Exo70 having a notably significant effect (Fig 2B). Consistent with these observations, we silenced each of the four subunits in MDBK cells and infected them with BoHV-1. Notably, the silencing of Exo70 led to the most substantial reduction in viral titer in the supernatant compared to the other three subunits (Fig 2C). At the same time, transmission electron microscopy showed that the number of virions released into the extracellular space was significantly lower in Exo70 knockdown (Exo70-KD) cells than in control cells (Fig 2D-2G), indicating that silencing Exo70 inhibited the extracellular release of herpesvirus. In summary, these results suggest that exocyst complex subunit deletion inhibits herpesvirus release and that silencing of Exo70 is particularly significant.

**Fig 2.**
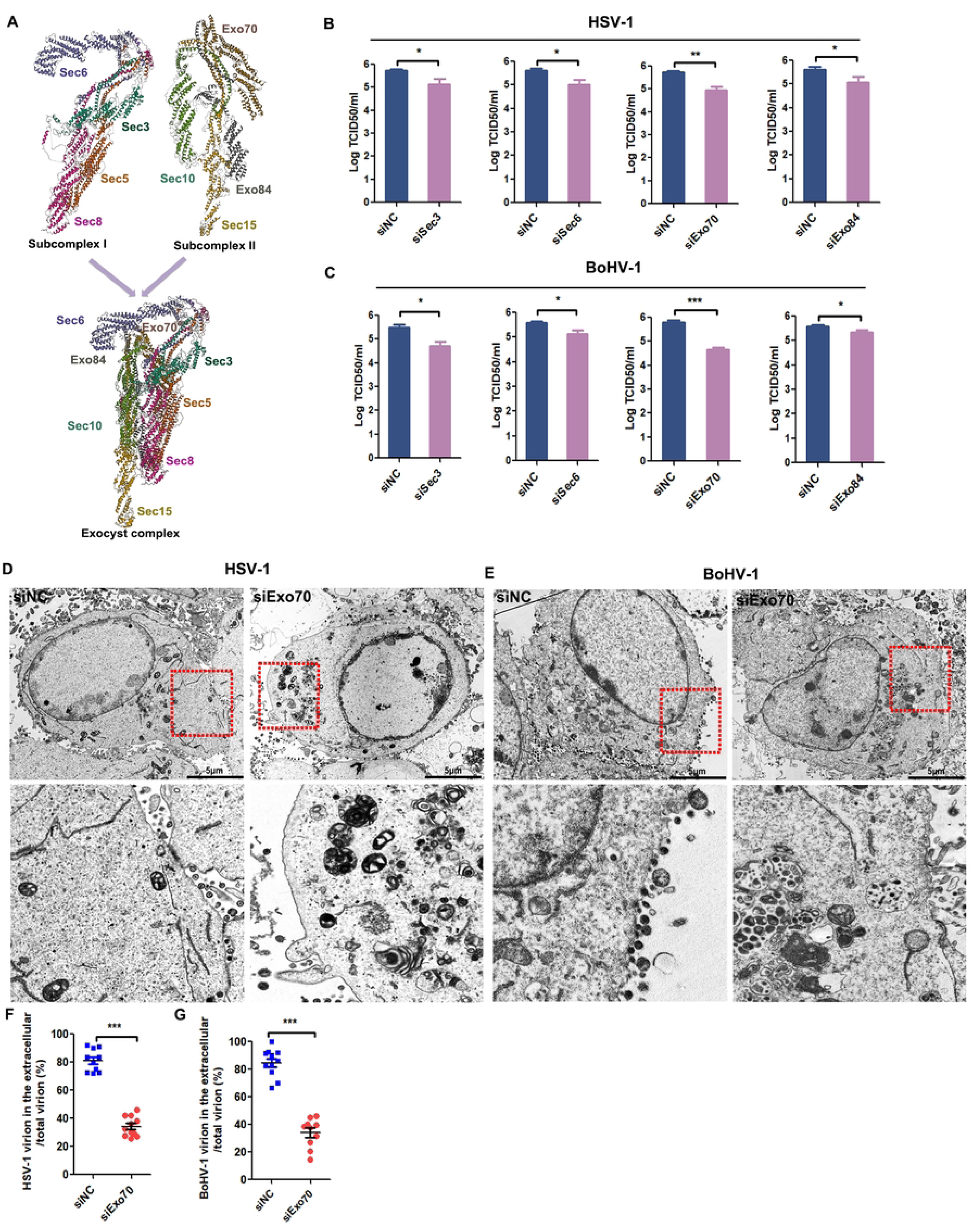
The deletion of exocyst complex subunits inhibits the herpesvirus replication. (A) Cryo-EM model of the exocyst complex (PDB: 5YFP). (B) and (C) MDBK and Vero cells transfected with siSec3, siSec6, shExo70, siExo84, or siNC were inoculated with HSV-1 or BoHV-1, and then the supernatant titers were determined by TCID50 assay. (D) and (E) Transmission electron microscopy showing the extracellular virions in WT or Exo70 knockdown cells infected with HSV-1 or BoHV-1. (F) and (G) The percentages of extracellular virions were quantified. Each dot represents one cell, and the median is represented by a line. Data are presented as the means ± SEMs of three independent experiments.

### Exo70 is vital for anchoring of viral secretory vesicles to the PM via its binding **to PI(4,5)P2**

Exo70, as a key subunit of the exocyst complex, participates in the intracellular transport of secretory vesicles, mediates the anchoring of secretory vesicles to the cell membrane, and promotes the excretion of substances in secretory vesicles [31]. To determine the impact of Exo70 on viral transport to the PM, we examined the localization of HSV-1 and BoHV-1 in Exo70-KD Vero cells and MDBK cells, respectively. We observed that in Exo70-KD cells, the gB protein accumulated in the cytoplasm, primarily localizing to the Golgi apparatus (Fig 3A, 3B, S2A, and S2B). Additionally, transmission electron microscopy revealed the presence of more virions near the Golgi apparatus in Exo70-KD cells than in control cells (Fig 3C and S2C). These data indicate that silencing of Exo70 blocked the intracellular transport of herpesvirus, leaving the virus stranded on the Golgi apparatus. In addition, immunofluorescence experiments demonstrated that the HSV-1 gB protein was dispersed in the cytoplasm and PM (labeled with Na^+^/K^+^ATPase) in siNC cells. However, in the siExo70 cells, the gB protein displayed cytoplasmic aggregation and limited distribution on the PM (Fig 3D). Consistently, Exo70 deficiency resulted in the BoHV-1 gB protein being mainly concentrated and distributed in the cytoplasm and rarely distributed on the cell membrane (Fig S2D). Notably, this phenomenon was similar to the effects of treatment with monensin. The above results indicated that Exo70 deficiency blocks the transport of herpesvirus to the PM.

**Fig 3.**
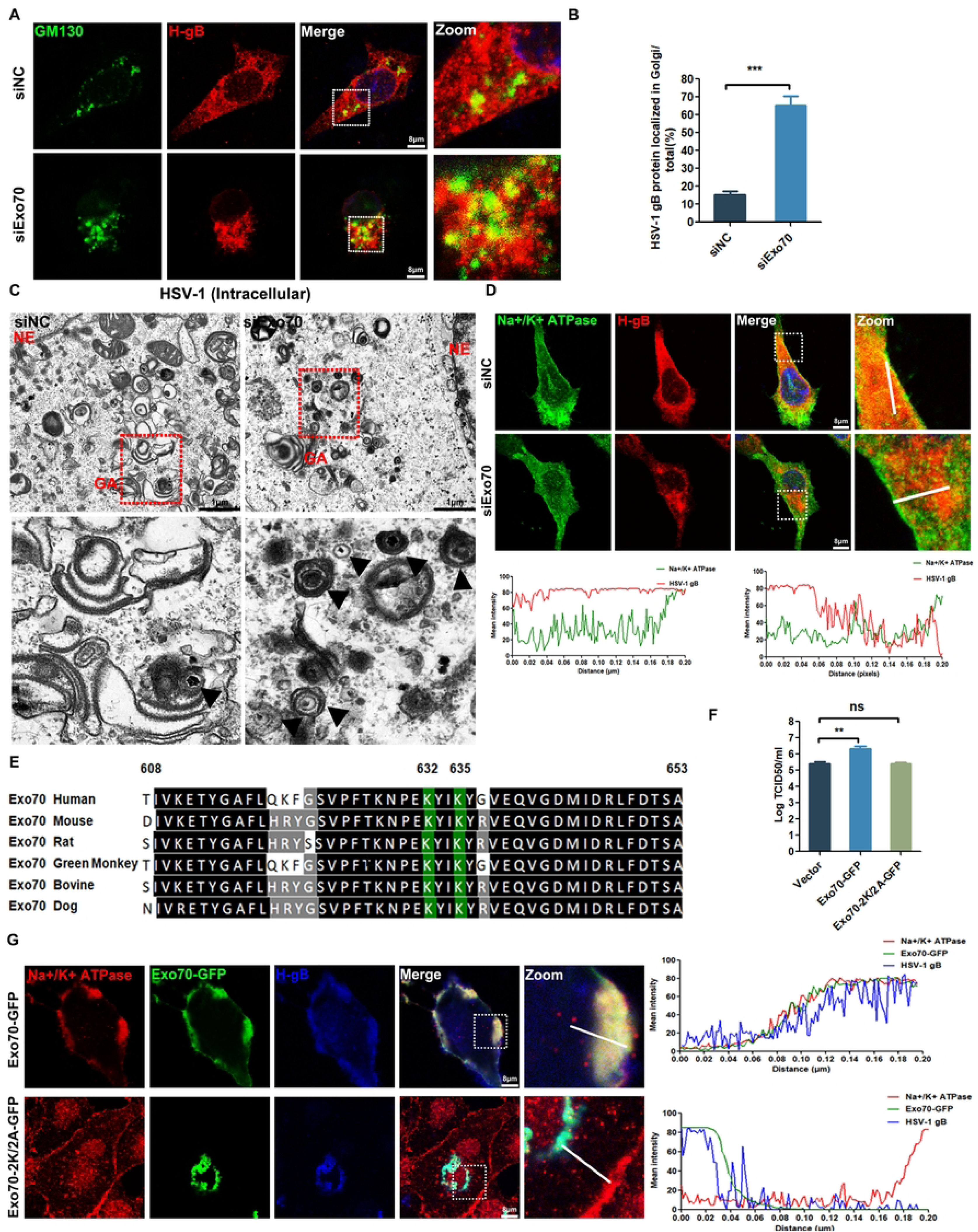
Knockdown of Exo70 limited the PM localization of the virus and caused the virus to remain on the Golgi apparatus. (A) Confocal microscope images of siNC and siExo70 Vero cells infected with HSV-1, fixed and stained for GM130 (green), gB (red), and chromatin (Hoechst, blue). (B) The percentages of gB protein localized in the Golgi were quantified. (C) Electron microscopy images of siNC and siExo70 Vero cells infected with HSV-1. Arrowheads indicate virions. NE, nucleus; GA,Golgi body, PM, plasma membrane. Scale bar, 1 μm. (D) Confocal microscopy images of siNC and siExo70 Vero cells infected with HSV-1, fixed and stained for Na^+^/K^+^ ATPase (green), gB (red), and chromatin (Hoechst, blue). The fluorescence intensity profiles of Na^+^/K^+^ ATPase (green) and gB (red) were measured along the line drawn by ImageJ. (E) Comparison of the amino acid sequences of Exo70 and its homologs. In variant residues are highlighted in white letters with gray boxes, and highly conserved residues are shown in white bolded text surrounded by black boxes. The residues K632 and K635 are marked with green boxes in the background. (F) Vero cells were transfected with the Exo70-WT-GFP, Exo70-2K/2A-GFP or Vector plasmid and infected with HSV-1 (MOI=0.1) for 24 h, after which the supernatant titer was determined by TCID_50_ assay. (G) Vero cells were transfected with plasmids encoding Exo70-WT-GFP or Exo70-2K/2A-GFP and then infected with HSV-1. Cells were fixed and stained for Exo70-GFP (green), Na^+^/K^+^ ATPase (red), and gB (blue). Scale bars: 8 μm. The fluorescence intensity profiles of Exo70 (green), Na^+^/K^+^ ATPase (red), and gB (blue) were measured along the line drawn by ImageJ. Data are presented as the means ± SEMs of three independent experiments.

The binding of Exo70 to PI(4,5)P2 is critical for the docking and fusion of post-Golgi secretory vesicles [18]. The C-terminal sequence of Exo70 is the most evolutionarily conserved region of this protein and a key domain for its direct binding to phospholipids. K632 and K635 are critical residues for the binding of Exo70 to PI(4,5)P2 [32]. We found that the K632 and K635 residues of Exo70 are conserved by comparing the C-terminal sequences of Exo70 in various mammals (Fig 3E). To assess the importance of K632 and K635 residues in the localization of Exo70 at the PM and anchoring of viral secretory vesicles, we generated a mutant Exo70-2K/2A, which residues K632 and K635 were substituted with alanine. Vero cells transfected with the GFP-tagged wild-type Exo70 (Exo70-WT) or Exo70-2K/2A plasmid were infected with HSV-1 and harvested at the indicated times. A TCID50 assay revealed an elevated virus titer in the supernatant of Exo70-WT cells, whereas the virus titer in the supernatant of Exo70-2K/2A cells was comparable to that of control cells (Fig 3F). Furthermore, immunofluorescence analysis showed that Exo70-WT colocalized with gB in the PM, whereas Exo70-2K/2A colocalized with gB in intracellular regions (Fig 3G), indicating that the residues K632 and K635 are required for the PM localization of HSV-1 gB. Overall, these findings strongly support the essential role of Exo70 in the anchoring of viral secretory vesicles to the PM.

### The Snapin-Exo70-SNAP23 axis facilitates fusion between viral secretory vesicles and the PM to promote virion egress

Following the tethering and docking of the vesicles on the PM, membrane fusion takes place. The Snapin-binding proteins Exo70 and SNAP23 are involved in GLUT4 vesicle docking and membrane fusion events [24]. We hypothesized that the Exo70-Snapin-SNAP23 axis is involved in the fusion of herpesvirus secretory vesicles with the PM. To test this hypothesis, we first detected the localization of the Exo70, SNAP23 and virus gB proteins in BoHV-1-infected cells. The results showed that the SNAP23, Exo70 and gB proteins colocalized in the PM after virus infection (Fig 4A). The amount of SNAP23 that was precipitated by Exo70 increased in a viral dose-dependent manner (Fig 4F). Subsequently, we examined the interactions between Snapin, Exo70 and SNAP23. The results indicated that Snapin interacted with Exo70 and SNAP23 (Fig 4B-4E). In addition, a reduction in Snapin by siSnapin transfection reduction the crosstalk between Exo70 and SNAP23 (Fig 4G), suggesting that Snapin aids in the assembly of the Exo70-SNAP23 complex during herpesvirus infection. Finally, we silenced the expression of SNAP23 in BoHV-1-infected cells that were overexpressing Exo70, and found that the knockdown of SNAP23 inhibited the virus release mediated by Exo70 (Fig 4H). All experimental data indicated that Snapin-Exo70-SNAP23 axis is involved in the membrane fusion and release of herpesvirus.

**Fig 4.**
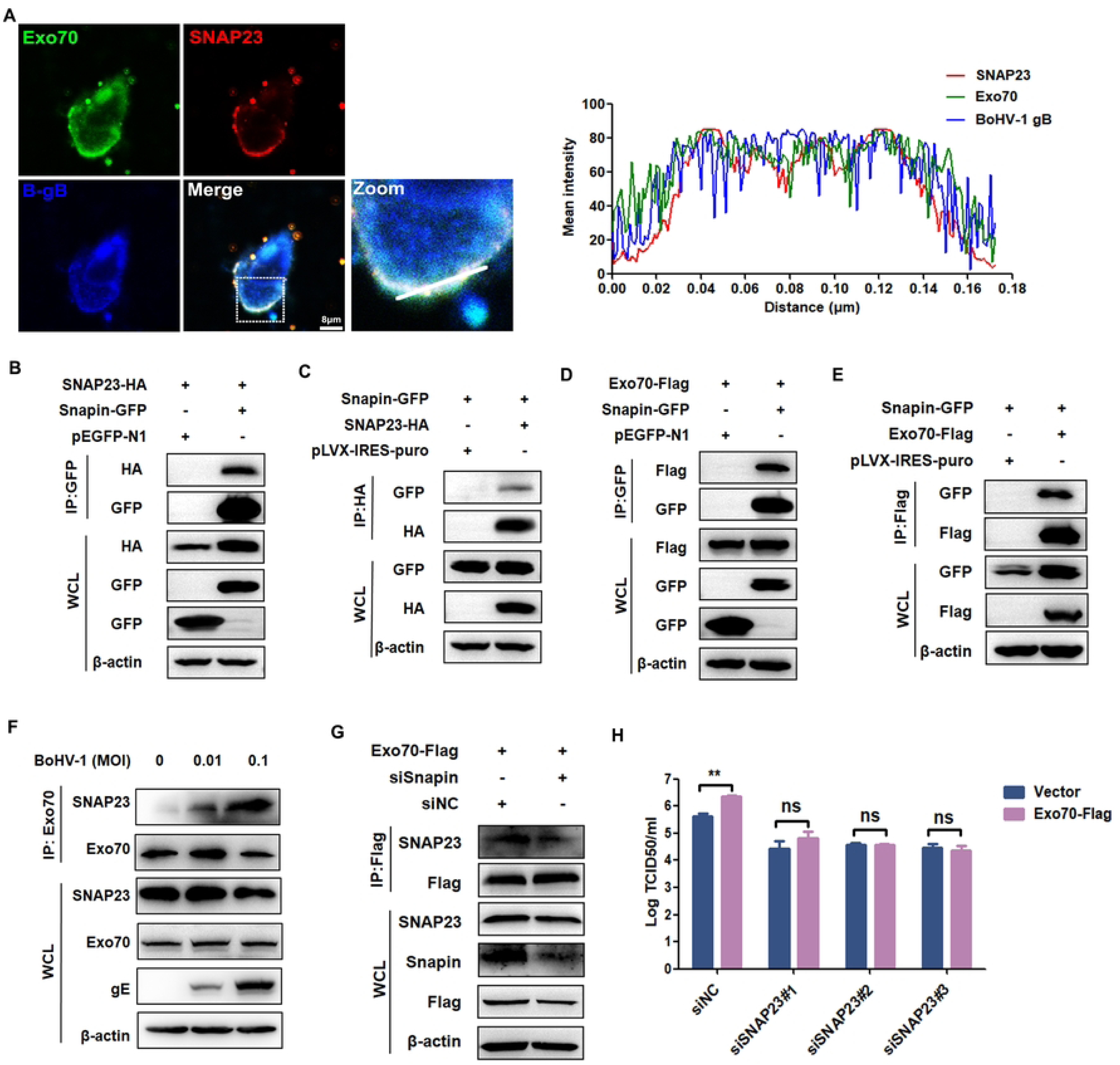
The Snapin-Exo70-SNAP23 axis is involved in viral secretory vesicle **membrane fusion and virion release.** (A) Immunofluorescence staining for Exo70, SNAP23, and gB in the BoHV-1-infected MDBK cells. Scale bars: 8 μm. (B-E) CoIP analyses the interaction between Snapin-GFP and SNAP23-HA or Exo70-Flag in HEK293T cells. (F) Detection of the association between Exo70 and SNAP23 by endogenous CoIP in MDBK cells that were infected with BoHV-1 at an MOI of 0, 0.01, and 0.1. (G) Exo70-Flag MDBK cells were transfected with siSnapin, CoIP analysis the interaction between Exo70 and SNAP23. (H) Exo70-Flag MDBK cells transfected with siSNAP23 and then infected with BoHV-1. The titer was determined by TCID_50_ assay. Data are presented as the means ± SEM of three independent experiments.

### Exo70 promoted virus release in an exocyst complex-dependent manner

Mutation of any exocyst subunit gene in yeast cells result in the aggregation of secretory vesicles and inhibition of cell secretion [33]. Negative stain electron microscopy (EM) of the purified complex also demonstrated a single compact octameric structure, with no visible subcomplexes [34]. Together, these data indicated that the exocyst complex exists as a single intact complex when it functions at sites of secretion. First, we selected BoHV-1 to infect MDBK cells to detect the interactions between excyst complex subunits. Exo70 antibodies were used to precipitate proteins in cells, and immunoblotting was used to detect the content of Sec6 in the sediments. The results showed that Exo70 binding to Sec6 increased with viral infection (Fig 5A). These results suggest that viral infection promotes the assembly of exocyst complex. In addition, immunoprecipitation experiments showed that the reduction of the Sec10 subunit by siSec10 transfection weakened the combination of Exo70 and Sec6 (Fig 5B). Confocal observation of viral localization showed that overexpression of Exo70 promoted the localization of the gB proteins to the PM, whereas the localization of gB to the PM was blocked when other subunits of the exocyst complex were absent (Fig 5C). Overexpression of Exo70 led to an increase in the quantity of virus in the supernatant but a decrease in the quantity of released virus was suppressed when individually silencing the other subunits of the complex (Fig 5D). All of the above data suggested that Exo70-mediated viral vesicle secretion events depend on the form of the exocyst complex.

**Fig 5.**
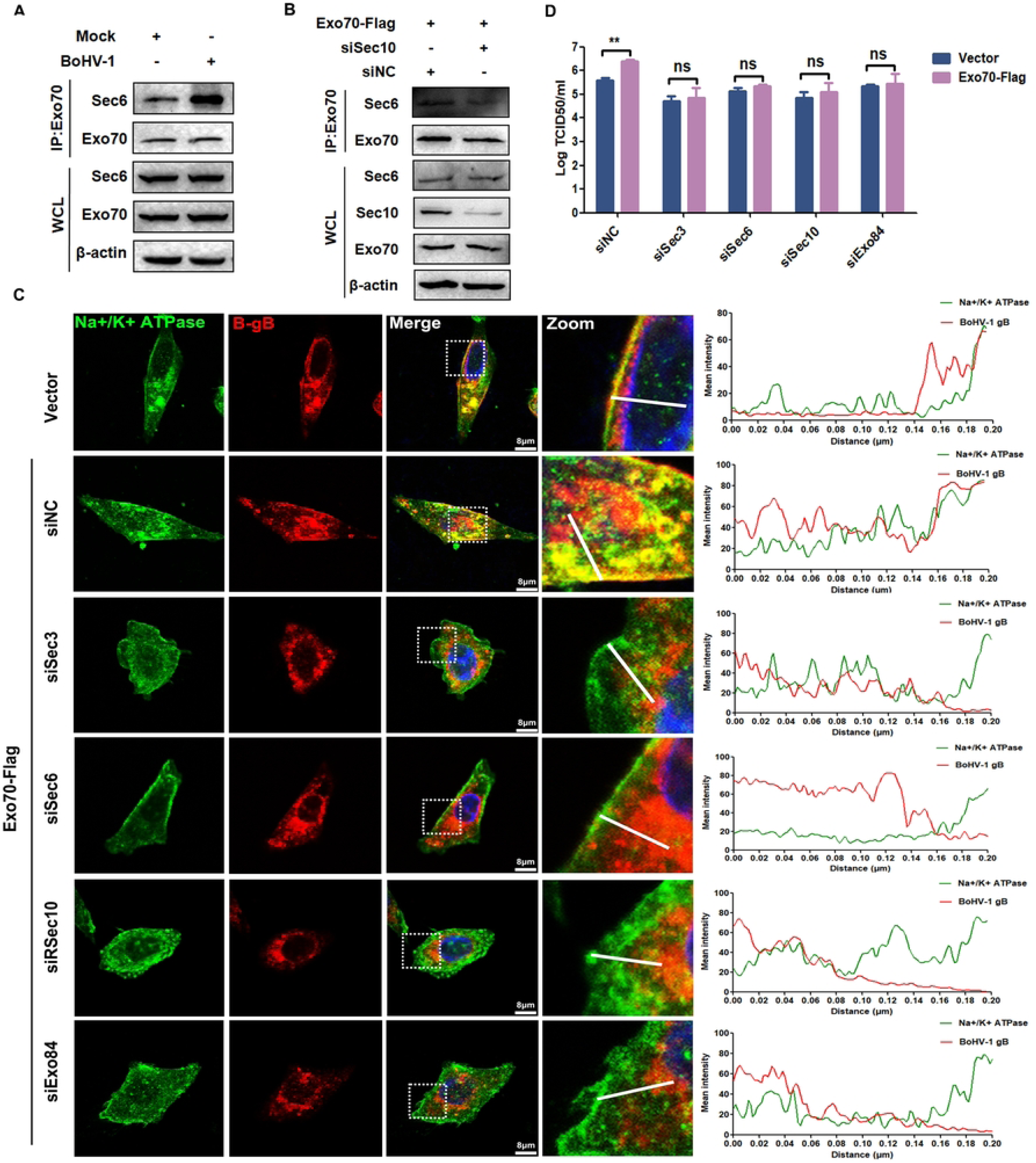
Exo70-mediated extracellular release of viruses depends on the form of the exocyst complex. (A) Detection of the association between Exo70 and Sec6 by endogenous co-immunoprecipitation and immunoblot analysis with the indicated antibodies in MDBK cells that were un-infected or infected with BoHV-1. (B) Exo70-Flag MDBK cells transfected with siSec10, CoIP analysis the interaction between Exo70 and Sec6. (C) and (D) Exo70-Flag MDBK cells transfected with siSec3, siSec6, siSec10, siExo84 or siNC and then infected with BoHV-1. The PM localization of BoHV-1 gB was evaluated by IFA. Scale bars: 8 μm. (C). The BoHV-1 titer was determined by TCID_50_ assay (D). Data are presented as the means ± SEM of three independent experiments.

### Rab11a is necessary for viral secretory vesicle trafficking mediated by Exo70 in an **exocyst complex-dependent manner**

Rab11 proteins have been demonstrated to have a crucial role in the transportation of viral proteins to the cell surface, including viruses like the Andes virus, HIV-1, Respiratory syncytial virus and MasonPfizer monkey virus [26, 35, 36]. Nevertheless, it remains unclear whether Rab11a has a role in the replication of BoHV-1 and the mechanism by which it promotes the transport of herpesvirus secretory vesicles. Initially, we infected MDBK cells with BoHV-1. At 12 h postinfection, we fixed the cells for immunofluorescence analysis using confocal microscopy. The results showed that Rab11a colocalized with the gB protein on the PM (Fig 6A). Moreover, our observations revealed the involvement of Rab11a in the intracellular transport of BoHV-1. We knocked down endogenous Rab11a in MDBK cells with siRNA, and at 12 h postinfection with BoHV-1, cells were detected by anti-gB antibody. We found that the gB protein was mainly concentrated in the Golgi apparatus and rarely distributed on the cell membrane in siRab11 cells (Fig 6B and 6C). Interestingly, this phenomenon resembled the effects observed upon knocked down of Exo70.

**Fig 6.**
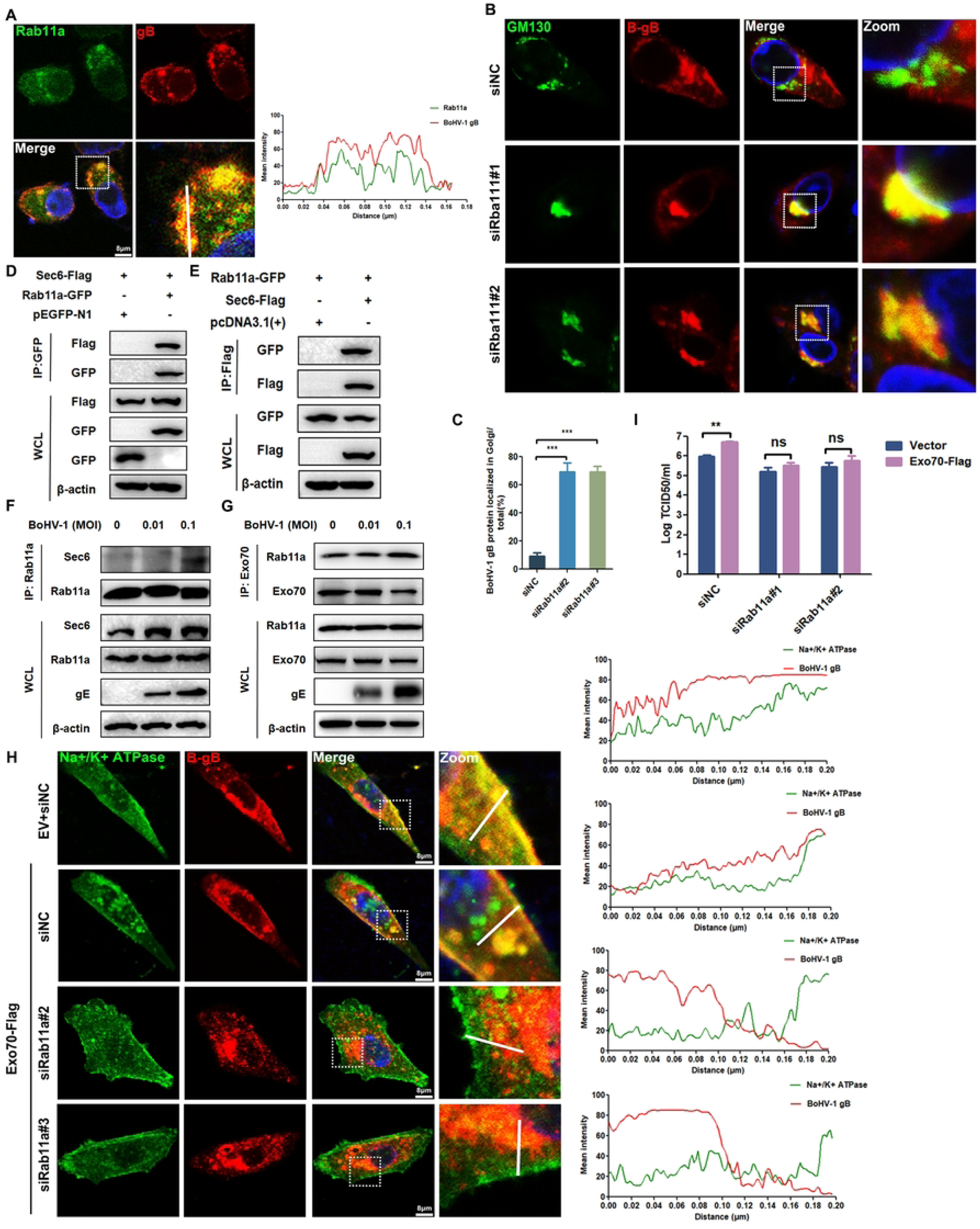
The exocyst complex mediates the intracellular transport of virions by binding Rab11a. (A) Immunofluorescence staining for Rab11a and gB in the BoHV-1-infected MDBK cells. (B) and (C) siRab11a cells and siNC cells were infected with BoHV-1. Immunofluorescence staining for stained for GM130 (green), gB (red), and chromatin (Hoechst, blue). (D) and (E) CoIP analyses the interaction between Rab11a-GFP and Sec6-Flag. (F) and (G) The association between Rab11a and Sec6 or Exo70 was detected by endogenous CoIP and immunoblot analysis with the indicated antibodies in MDBK cells that were uninfected (left) or infected with BoHV-1 at MOI of 0.01, and 0.1. (H) Exo70-Flag MDBK cells transfected with siRab11a and then infected with BoHV-1. The PM localization of BoHV-1 gB was evaluated by IFA. Scale bars: 8 μm. The fluorescence intensity profiles of Na^+^/K^+^ ATPase (green) and gB (red) were measured along the line drawn by ImageJ. (I) The BoHV-1 titer was determined by TCID_50_ assay. Data are presented as the means ± SEM of three independent experiments.

Next, we investigated the involvement of Rab11a in the intracellular transport of virus particles mediated by the exocyst complex. HEK293T cells were transfected with GFP-tagged Rab11a alone or in combination with Flag-tagged Sec6. The immunoprecipitation assay yielded results showed that Rab11a could bind the exocyst complex subunit Sec6 (Fig 6D and 6E). To explore the interaction among the virus, Rab11a and the exocyst complex, cells were infected with different viral doses, and binding was detected. The data indicated that the precipitation of Sec6 by Rab11a and that of Rab11a by Exo70 increased in a viral dose-dependent manner (Fig 6F and 6G). These findings indicate that viral infection enhances the interaction between Rab11a and the exocyst complex. In addition, knockdown of Rab11a by siRab11a inhibited the localization of the gB protein on the PM and suppressed the virus release mediated by Exo70 (Fig 6H and 6I). In brief, these results indicate that Rab11a is necessary for exocyst complex-mediated intracellular transport.

### ES2, an inhibitor of exocytosis mediated by exocyst complex, suppressed the release of herpesvirus and alleviated the symptoms of HSV-1 infection in BALB/c mice

Endosidin2 (ES2) was identified through plant-based chemical screening as a transport inhibitor that combines with the Exo70 subunit to terminate the final stage of exocytosis [17, 37]. ES2 treatment can effectively improve the cisplatin sensitivity of acquired cisplatin resistant epithelial ovarian cancer cells, and can be used as a potential drug for the treatment of epithelial ovarian cancer [38]. In addition, ES2 treatment can lead to impaired transport of GLUT4 to the PM and hinder glucose uptake under insulin stimulation [39]. ES2 has the potential to be a treatment for human diseases caused by dysfunction of the exocyst complex. Whether ES2 regulates the process of herpesvirus infection mediated by the exocyst complex is unclear. Therefore, we next explored the impact of the Exo70-targeting drug ES2 on the transport and release of herpesvirus. First, in virus-infected cells treated with ES2, the localization of Exo70 on the PM was blocked, while the gB protein was predominantly located in the cytoplasm, and its localization on the cell membrane was limited (Fig 7A, 7B, S3A, and S3B), which is consistent with the results observed following Exo70-KD. Additionally, treatment with ES2 did not impact Exo70 expression (Fig 7C), but inhibited virus release compared to the DMSO-treated group (Fig 7D and S3C). The above results indicated that indicated that ES2 blocked the release of herpes virus by suppressing Exo70 localization on the PM.

**Fig 7.**
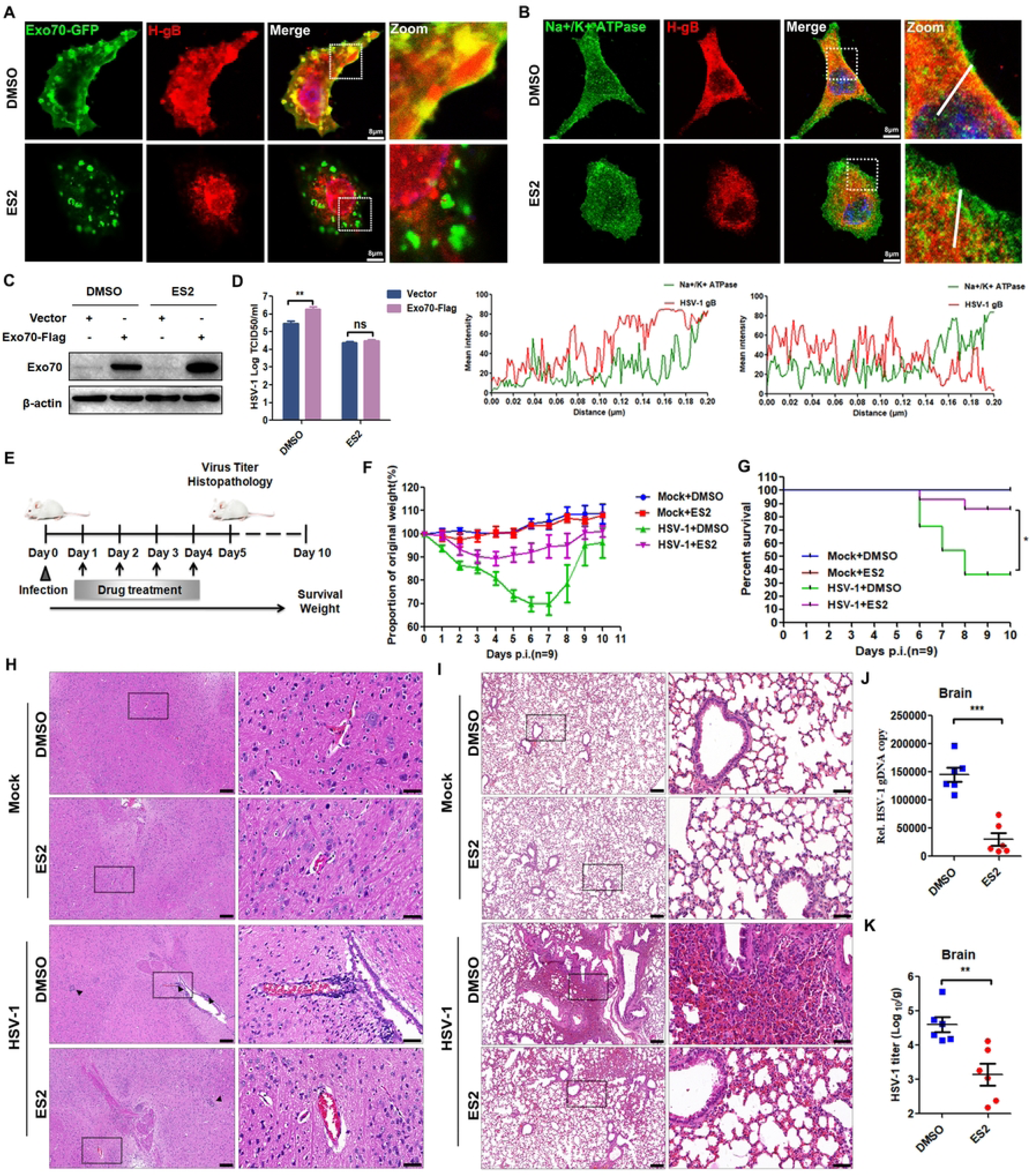
ES2 could protect cells and mice from α-herpesvirus infection by targeting Exo70. (A) Vero cells were treated with DMSO or ES2 and infected with HSV-1. Immunofluorescence staining for Exo70 and gB. Scale bars: 8 μm. (B) Confocal microscopy images of Vero cells treated with DMSO or ES2, infected with HSV-1, fixed and stained for Na^+^/K^+^ ATPase and gB. (C) and (D) Exo70-Flag Vero cells were treated with DMSO or ES2 and inoculated with HSV-1. Then, the expression of Exo70 protein was assayed by western blotting, and the supernatant titers were determined by TCID_50_ assay. (E) The ES2 treatment regimen applied to BALB/c mice infected with HSV-1 from 0 to 10 days. (F) and (G) Age- and sex-matched BALB/c mice (7 weeks of age, male) were infected with HSV-1 (1×10^5^ pfu/mouse) or DMEM and then intraperitoneally injected with DMSO or ES2 (10 mg/kg), and changes in the survival rate and body weights of the mice were recorded. (H) and (I) Hematoxylin and eosin (HE) staining of brain and lung sections of mice treated as described in (E) followed by intracerebral injection of HSV-1 for 5 days. Scale bar, 50 μm. (J) and (K) qRT‒PCR analysis of HSV-1 genomic DNA in the brains of mice. TCID_50_ assays analyzing HSV-1 titers in the brains of mice. Data are presented as the means ± SEM of three independent experiments.

Next, we examined the physiological function of ES2 in resisting HSV-1 infection in mice. BALB/c mice were injected with HSV-1 or DMEM as a control. The next day, the mice were intraperitoneally injected with ES2 or DMSO as a control. The mice underwent continuous intraperitoneal injection for 4 days, and physiological changes were continuously monitored for 10 days (Fig 7E). As shown in Figure 7F and 7G, ES2 had no effect on body weight or survival rate in the uninfected mice, while the mice infected with HSV-1 showed less weight loss after intraperitoneal injection of ES2, and their survival rate was significantly higher than that of the mice injected with DMSO, indicating that ES2 enhanced the ability of mice to resist HSV-1 infection. After 5 days of HSV-1 infection, the brain and lung of the mice were taken for histopathological examination. Hematoxylin and eosin staining revealed that the brain and lung lesions in ES2-injected mice were less severe than those in the DMSO-injected mice. There was a noticeable reduction in inflammatory cell infiltration and tissue damage in the ES2-treated group (Fig 7H and 7I). Additionally, the relative copy number of HSV-1 genomic DNA (gDNA) in the brains of ES2-injected mice was lower compared to that in the DMSO-injected mice, indicating a subsequent decrease in HSV-1 replication (Fig 7J). Consistent with the above result, the titer of HSV-1 in the brains of the ES2-injected mice was lower than that in the brains of DMSO-injected mice (Fig 7K). In summary, the above results indicate that ES2 enhanced the resistance of mice to HSV-1 infection and alleviated pathological symptoms in the infected mice.

## Discussion

The exocyst complex is an evolutionarily conserved, multisubunit protein complex involved in tethering secretory vesicles to the PM prior to fusion mediated by SNAREs [40]. However, it remains unclear whether the exocyst complex is also involved in tethering virus secretory vesicles to the PM. Mature herpesvirus particles are released from cells through a fusion mechanism similar to exocytosis, wherein vesicles containing mature virus fuse with the PM [41, 42]. Our study revealed that deleting exocyst complex subunits reduced the viral load in the supernatant, suggesting the involvement of the exocyst complex in viral replication. Notably, the knockdown of Exo70 had a particularly significant effect. In addition, in Exo70-KD cells, the number of extracellular virions decreased while the number of intracellular virions increased, indicating that knockdown of Exo70 inhibited the extracellular release of the virus. Although Exo70 participates in the intracellular transport of secretory vesicles generated at the Golgi, mediates the anchoring of secretory vesicles on the cell membrane by directly binding PI(4,5)P2, and promotes the fusion of secretory vesicles with the PM [43, 44], it is not clear whether Exo70 is involved in herpesvirus vesicle anchoring to the PM. In HSV-1 or BoHV-1 infected cells, the viral gB protein was distributed in the cytoplasm and PM in control cells, whereas in Exo70-KD cells, the gB protein accumulated in the cytoplasm, primarily localizing to the Golgi apparatus. The above results showed that Exo70 is involved in the PM localization of herpesviruses. Exo70 associates with the PM by utilizing a surface patch of basic residues at its C-terminus that interacts with PI(4,5)P2 [18]. Blocking the localization of Exo70 at the PM was achieved by substituting endogenous Exo70 with mutants lacking the ability to bind PI(4,5)P2. This restriction of Exo70 localization also led to limitations in the localization of the virion at the PM, highlighting the necessity of Exo70 localization for the anchoring of the virion to the PM.

Fusion is the final step of vesicle release and is mediated by a family of proteins called SNAREs [21]. v-SNAREs specifically pair with their cognate t-SNAREs, forming the “SNARE complex” that facilitates fusion between the vesicle and target membranes [21]. Although the t-SNARE protein SNAP23 is interacts with Exo70 [45], but the effects of this interaction on virus release remain unknown. In this study, we found that Exo70, SNAP23 and virions colocalized at the PM and that viral infection promoted the interaction between Exo70 and SNAP23. In addition, the absence of SNAP23 inhibited Exo70-mediated virus release, indicating that SNAP23 plays a role in the process by which Exo70 promotes virion egress. The interaction between Exo70 and SNAP23 is mediated by Snapin in GLUT4 vesicle trafficking [24]. However, it remains unclear if Snapin also promotes this interaction during virus infection. Our data show that Snapin silencing attenuated the interaction between Exo70 and SNAP23 during BoHV-1 infection. The aforementioned findings suggest that the Exo70-Snapin-SNAP23 axis significantly contributes to the fusion of herpesvirus secretory vesicles with the cell membrane.

An intact exocyst complex is necessary for the completion of exocytosis [46]. Inhibiting the expression of Sec6, Sec8, and Exo70 subunits through transfection with siRNA resulted in the inhibition of GLUT4 exocytosis to the PM in adipocytes [47, 48]. The study aimed to investigate whether the intact exocyst complex is responsible for the release of herpesvirus mediated by Exo70. Since the stability of the exocyst complex depends on the presence of all subunits, we disrupted the integrity of the complex by silencing Sec3, Sec6, Sec10, and Exo84 with siRNA. Our data showed that the destruction of the exocyst complex blocked Exo70-mediated PM localization and the extracellular release of virions, indicating that Exo70 regulates the extracellular release of viruses in a manner dependent on the integrity of the exocyst complex.

The exocyst complex relies on the Rab protein Rab11 to facilitate vesicle transport to the PM for exocytosis. Rab11 is a crucial regulator of membrane trafficking from the TGN and the recycling of endosomes to the PM [49, 50]. The exocyst complex component Sec6 recruits and activates Rab11a, facilitating the expulsion of intracellular uropathogenic *E. coli* from their intracellular niche [27]. Our data demonstrates that Rab11a has the ability to interact with Sec6. It is possible that Rab11a is involved in the assembly and transport of herpesvirus vesicles. Rab11a is present on the pseudorabies virus secretory vesicles, and is involved in the formation of endocytic tubules for the HSV-1 secondary envelope [51, 52]. In this study, we found that Rab11a colocalizes with the gB protein during BoHV-1 infection, indicating that Rab11a may be involved in the intracellular transport of BoHV-1. Further research confirmed the above hypothesis. Knockdown of Rab11a prevented the virus from being transported to the PM, causing it to be retained in the Golgi apparatus. Rab11a interacts with both the exocyst subunit and herpesvirus glycoprotein, with viral infection enhancing its interaction with the exocyst complex. Furthermore, knockdown of Rab11a reduced the PM localization and extracellular release of virions, which were mediated by Exo70. This suggests that Rab11a acts as a “bridge” between herpesvirus secretory vesicles and the exocyst complex.

The small molecule ES2 was identified as an inhibitor of Exo70. In studies of vesicular transport in *Arabidopsis thaliana*, ES2 was found to inhibit the localization of Exo70 to the PM by binding L598 and I613, resulting in the inhibition of exocytosis and endosomal recycling in both plant and human cells and the enhancement of plant vacuolar trafficking [17]. ES2 was further proven to overcome acquired cisplatin resistance in epithelial ovarian cancer cells in vitro and in vivo and could be further developed as a lead compound to improve the clinical chemotherapeutic efficacy of cisplatin in epithelial ovarian cancer [38]. Furthermore, treatment of skeletal myoblast cells with ES2 impairs GLUT4 trafficking to the PM and hinders glucose uptake in response to insulin stimulation, suggesting the potential utility of ES2 as a tool for studying resistance in type 2 diabetes [39]. Therefore, ES2 is expected to become a target drug for the treatment of human diseases caused by exocyst complex dysfunction, such as diabetes and cancer. However, whether ES2 can be used as an antiviral agent needs to be further evaluated. Therefore, in this study, we evaluated the antiviral effect of ES2 through in vitro and in vivo experiments. In virus-infected cells, ES2 treatment restricts the localization of Exo70 at the PM without affecting its expression, as demonstrated by in vitro experimental data. Additionally, the gB protein of the virus primarily localizes in the cytoplasm, its PM localization is blocked by ES2, and this leads to a decrease in the viral titer in the supernatant, indicating the ability of ES2 to inhibit extracellular herpesvirus release by restricting the localization of Exo70 to the PM. In vivo experiments demonstrated that intraperitoneal injection of ES2 at a dose of 10 mg/kg improved the weight and survival rate of HSV-1-infected mice, alleviated their pathological symptoms, and reduced the viral load, indicating the potential efficacy of ES2 in treating HSV-1 infection. In summary, all data indicate that ES2, as an inhibitor of Exo70, has potential as a treatment for herpesvirus infection.

In this study, we propose a model in which the exocyst complex is involved in the transport, anchoring and release of viruses (Fig 8). Following the formation of the secondary envelope of the herpesvirus, the exocyst complex binds to GTPase Rab11a on vesicles containing virions and transports the virions to the PM. Then, the exocyst complex anchors the virions to the PM through its subunit Exo70, which interacts with PI(4,5)P2. Finally, under the action of Snapin, Exo70 interacts with SNAP23, leading to the fusion of virions with the PM and eventual release into the extracellular space. More importantly, we propose the potential of ES2 as an antiviral drug that targets Exo70.

**Fig 8.**
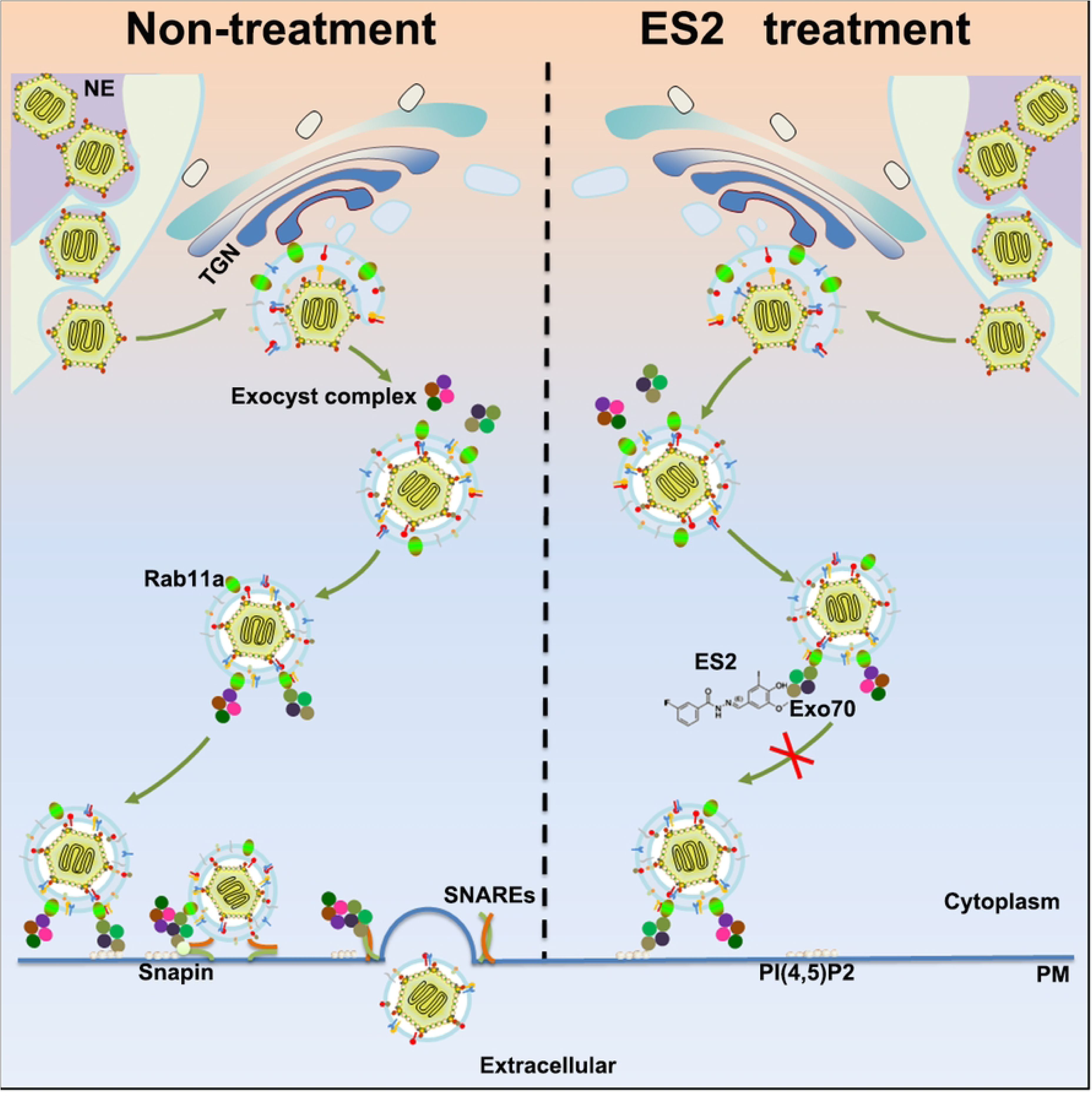
Pattern diagram of exocyst complex-mediated α-herpesvirus particle anchoring, fusion, and release.

## Materials and methods

### Ethics statement

All animal studies were approved by the Institutional Animal Care and Use Committee of Shandong Normal University. Studies were conducted under animal biosafety level 2 (ABSL-2) containment and following the Guide for the Care and Use of Agricultural Animals in Research and Teaching.

### Cells and viruses

HEK293T, Vero E6, and Madin-Darby Bovine Kidney (MDBK) cells were cultured in Dulbecco’s modified Eagle’s medium (DMEM; BI) supplemented with 10% fetal bovine serum (FBS; HyClone), penicillin and streptomycin (100 U/ml; New Cell & Molecular Biotech Co., Ltd). Cells were maintained at 37°C in a 5% CO_2_ environment. The BoHV-1/BarthaNu/67 strain (HVRIIBRV0004) was obtained from the China Veterinary Culture Collection Center (CVCC). Herpes simplex virus 1 (ATCC-2011-9 strain; VR-1789) was obtained from American Type Culture Collection (ATCC).

### Mouse experiments

Specific-pathogen-free BALB/c mice were purchased from Jinan Pengyue Experimental Animal Breeding Co., Ltd. For infection experiments, age- and sex-matched BALB/c mice (7 weeks of age) were anesthetized with sodium pentobarbital (30-50 mg/kg body weight), inoculated intracranially with HSV-1 (1×10^5^ pfu/mouse) or DMEM, and then intraperitoneally injected with DMSO or ES2(10 mg/kg). At the indicated times postinfection, the mice were euthanized, and the brain was harvested for histological analyses. Survival, weight loss, and disease signs were monitored for 10 days.

### Electroporation and establishment of stable cell lines

MDBK cells were transfected with either Exo70-Flag plasmid or an empty vector (EV) using electroporation. After 48 hours, puromycin was added to the cells at a concentration of 2.2 μg/ml. The cells were then screened for 4 days until drug-resistant cell clones were obtained. From these clones, drug-resistant cells were selected and the culture was expanded. Western blot analysis was performed to assess the expression of Exo70 in MDBK cells with stable expression of the corresponding genes or EV cells.

### RNAi

Cells were grown to 40% confluence and then incubated with 50 nM siRNA and Attractene transfection reagent (QIAGEN) for 36 h according to the manufacturer’s instructions.

### Immunofluorescence confocal microscopy

MDBK or Vero cells in the logarithmic growth phase were suspended in a cell suspension and seeded onto 24-well plates with coverslips. When the cells reached 80% confluence on the coverslips, the original culture medium was discarded, and the cells were washed with PBS, fixed with room temperature 4% paraformaldehyde (PFA)/PBS for 30 min, washed, permeabilized with 0.1% Triton X-100/TBS for 30 min, and blocked with 0.5% BSA. Next, they were washed and sequentially incubated with primary antibody at 4°C overnight and secondary antibody at room temperature for 60 min. Nuclei were stained with Hoechst (Solarbio). Images were acquired using a LEICADMI8 confocal microscope.

### Electron microscopy

The virus-infected or plasmid-transfected cells were harvested by cell scraping, centrifuged at 3000 rpm for 2 min, fixed with 2.5% glutaraldehyde at room temperature for 2 h, and then transferred to 4°C. Then, the cells were dehydrated with sequential washes in 50%, 70%, 90%, 95%, and 100% ethanol. Areas containing cells were block mounted and thinly sliced. The sections were photographed using a Hitachi HT7800 transmission electron microscope (Hitachi), and the images were processed with the Hitachi TEM system.

### Immunoblot analysis and coimmunoprecipitation

For immunoblot analysis, cells were lysed with RIPA lysis buffer (PMSF and a complete protease inhibitor cocktail (MCE)) on ice for 30 min. A coimmunoprecipitation assay was performed using the Pierce^TM^ Co-IP kit (Thermo Scientific, Waltham, MA, USA) according to the manufacturer’s protocol. Whole-cell extracts were collected 48 h after transfection and lysed in lysis buffer supplemented with 1 mM PMSF and a complete protease inhibitor cocktail on ice for 30 min. After centrifugation for 10 min at 13,000×g and 4°C, the supernatants were collected and incubated with Protein-G Sepharose beads coupled to specific antibodies for 2 h or overnight with rotation at 4°C, and the beads were washed 3 times with lysis buffer. Bound proteins were eluted by boiling for 5 min with sample buffer (50 mM Tris-HCl (pH=6.8), 2% SDS, 10% glycerol, 0.1% bromophenol blue and 1% β-mercaptoethanol). The lysates were subjected to immunoblotting. For immunoblot analysis, immunoprecipitates or whole-cell lysates were separated by SDS–PAGE, electro-transferred to PVDF membranes and blocked for 1 h with a 5% nonfat milk solution, followed by blotting with the indicated antibodies and detection by an Omni-ECL™ Femto Light Chemiluminescence Kit (Epizyme).

### TCID50 assay

TCID50 was determined in Vero or MDBK cells infected with 10-fold serially diluted viruses and cultured at 37°C in a 5% CO2 atmosphere for 1 h, the cells were washed three times and incubated in serum-free DMEM. Supernatants were collected at 24 hpi and titrated on Vero or MDBK cells. TCID50 value was calculated using the Reed-Muench method. Each experiment was conducted three times, and each experiment was performed in triplicate.

### Histopathological analysis

Brains and lungs from mock-infected or HSV-1-infected mice were dissected, fixed in 4% paraformaldehyde, dehydrated, embedded in paraffin, sectioned, stained with a hematoxylin and eosin (H&E) solution, and then examined by microscopy (3D HISTECH) for histological changes.

### Quantification and statistical analysis

The results are expressed as the means ± SEMs. All graphs were generated with GraphPad Prism 5.0. Values of *p < 0.05 were considered statistically significant.

## AUTHOR CONTRIBUTIONS

HBH and HMW developed the concept of the study; HBH and WQM designed the experiments; WQM performed the experiments; JLX participated in the virus infection experiments. JY was involved with the IF assay. YNX and LW were involved with the IP assay. YNX helped construct the recombinant plasmids. LTH and FCS helped with the mouse experiments. WQM and DH collected and analyzed the data; WQM wrote the manuscript. All the authors read and approved the final manuscript.

## ACKNOWLEDGMENTS

This work was partially supported by grants from the National Natural Science Fund of China (32202772 and 32072834), Taishan Scholar and Distinguished Experts (H. H, tspd20181207), Jinan Innovation Team (202228060).

